# Ghrelin-Reactive Autoantibodies are elevated in Children with Prader-Willi Syndrome

**DOI:** 10.1101/093930

**Authors:** Gabrielle Crisp, Ohn Nyunt, Lisa Chopin, Inge Seim, Mark Harris, Penny Jeffery

**Affiliations:** Ghrelin Research Group, Translational Research Institute-Institute of Health and Biomedical Innovation, School of Biomedical Sciences, Queensland University of Technology, Woolloongabba, Queensland, Australia; Royal North Shore Hospital, New South Wales, Australia; Lady Cilento Children’s Hospital, South Brisbane, Queensland, Australia; Mater Research Institute – University of Queensland, Translational Research Institute, Woolloongabba, Queensland, Australia

## Abstract

Prader-Willi Syndrome (PWS) is a complex genetic disorder characterized by developmental and growth abnormalities, insatiable appetite, and excessive eating (hyperphagia). The underlying cause of hyperphagia in PWS is currently unknown, however, elevated levels of the peptide hormone ghrelin is believed to contribute. Recently, ghrelin-reactive autoantibodies (isotype IgG) were identified in non-genetic obesity. These autoantibodies act as ghrelin carrier proteins and potentiate its orexigenic effects. Here, we describe the identification of ghrelin-reactive autoantibodies in a cohort of 16 children with PWS. In comparison to unaffected siblings, autoantibody levels are significantly increased in PWS children. We further show that autoantibody levels are unaffected by food intake, unlike plasma ghrelin which declines postprandially in both groups. Critically, we also demonstrate that the autoantibodies bind the major circulating ghrelin isoforms, unacylated ghrelin, which does not stimulate appetite, and the orexigen acylated ghrelin. In excess, unacylated ghrelin may compete with acylated ghrelin for autoantibody binding. Taken together, this is the first report on ghrelin-reactive antibodies in a pediatric population, and the first to demonstrate that the antibodies do not discriminate between orexigenic and non-orexigenic ghrelin isoforms. Our work suggests that ghrelin autoantibodies can be targeted using non-orexigenic forms of ghrelin, thereby providing a novel therapeutic target for PWS and for obesity in general.

## Introduction

Prader-Willi Syndrome (PWS) is the most common genetic cause of obesity in children and is characterized by developmental and growth abnormalities, insatiable appetite and impaired satiety [1]. PWS patients also exhibit high levels of the orexigenic peptide hormone ghrelin. The two major forms of ghrelin in the circulation are acyl ghrelin, which potently stimulates appetite and food-seeking behaviour, and unacylated ghrelin (UAG), which has no effect on appetite [2–4]. Recently, ghrelin-reactive autoantibodies (isotype IgG) were identified in non-genetic obesity [5]. These autoantibodies bind ghrelin reversibly, acting as carrier proteins that protect ghrelin from degradation and potentiate its orexigenic effects [5]. Here, we sought to further explore this association by characterizing ghrelin autoantibodies in children with PWS and non-affected sibling controls.

## Methods

Sixteen children with PWS and 16 controls, matched for body mass index (BMI), were recruited to the study. Plasma was collected after an overnight fast (baseline), and 10, 20, 30, 60 and 120 minutes after a standardized mixed meal. Plasma acylated ghrelin levels were measured by ELISA (Human Active Ghrelin ELISA, EZGRA-88K, Millipore). Ghrelin-reactive IgG levels were measured using an adapted ELISA method [6]. To test specificity, and to determine if the autoantibodies also bind unacylated ghrelin (UAG), samples were pre-absorbed overnight with 10^−6^ M synthetic acylated ghrelin or UAG (Mimotopes, Australia) prior to the ELISA.

## Results

PWS children were shorter in stature and displayed reduced lean mass compared to controls (Table 1). Mean fasting plasma acylated ghrelin levels were significantly higher in the PWS group (*P* < 0.01, Bonferonni-corrected two-way ANOVA; Figure 1A and Table 1), but postprandially ghrelin levels were similar to those in the control group after 60 minutes. Unlike acylated ghrelin, postprandial levels of ghrelin-reactive autoantibodies remained unchanged in both PWS and control children (*P* > 0.05; Figure 1B). Children with PWS exhibited significantly higher fasting and postprandial levels of plasma ghrelin-reactive autoantibodies than sibling controls (fasting comparison *P* < 0.0001, unpaired Student’s t-test; Figure 1B). Pre-absorption of plasma with either acylated ghrelin or the non-orexigenic isoform, UAG, decreased autoantibody levels in the PWS and control groups (*P* < 0.001, unpaired Student’s t-test; Figure 1C, D).

**Figure 1:**
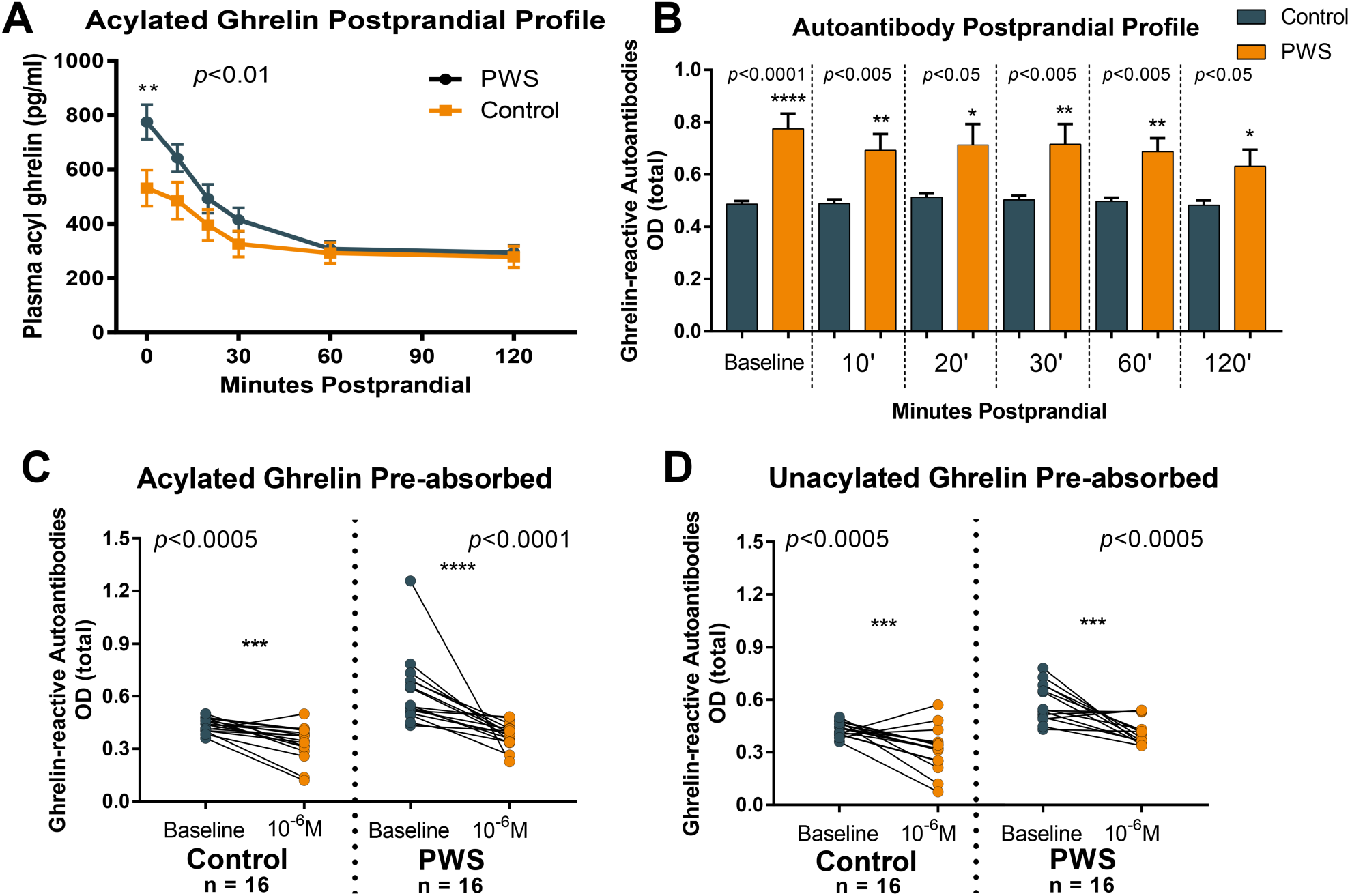
A) Plasma acylated ghrelin levels. B) Ghrelin-reactive autoantibodies are increased in PWS across the entire postprandial profile in the PWS group. C) Acylated ghrelin pre-absorption. D) Unacylated ghrelin (UAG) pre-absorption.

**Table 1:**
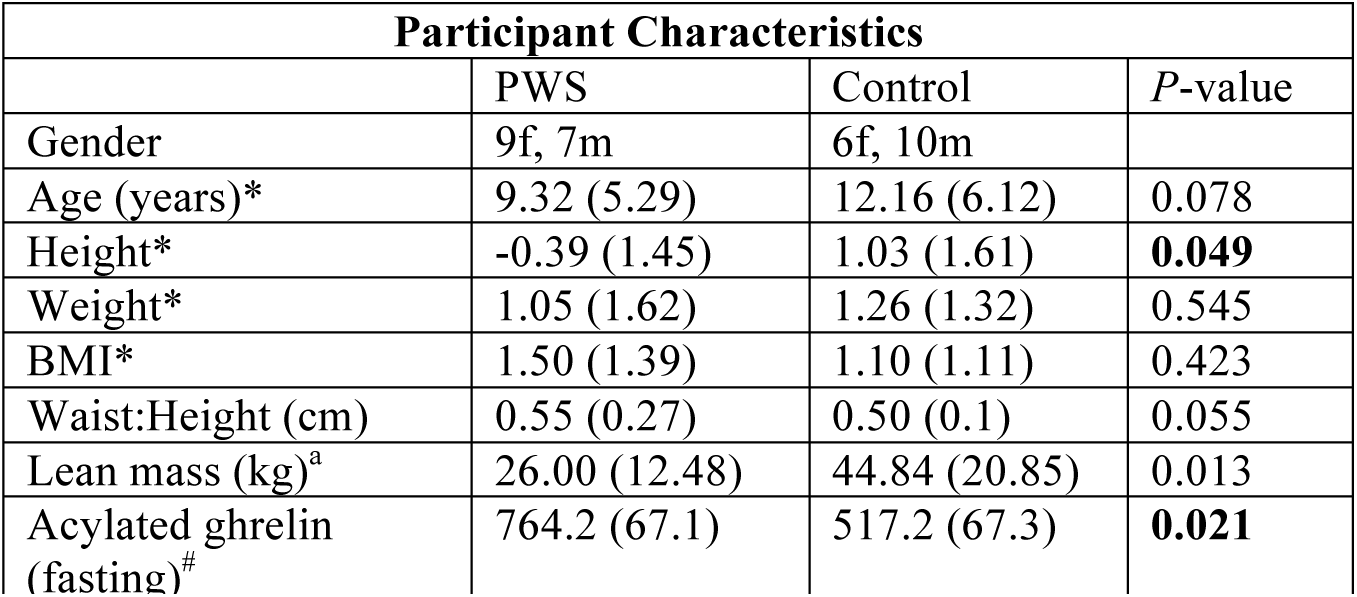
Participant characteristics. Abbreviations: IgG, Immunoglobulin G; PWS, Prader-Willi Syndrome; f, female; m, male; BMI, body mass index. *Standard Deviation Score (Median; Interquartile range); cm, centimetres; ^a^Mean (± S.D.); #Mean (± S.E.M; pg/mL). Difference between PWS and control participants was determined using Student’s t-test.

## Discussion

To our knowledge, this study represents the first evidence of an association between Prader-Willi Syndrome and ghrelin-reactive autoantibodies. Autoantibody levels were higher in children with PWS compared to sibling controls, both after fasting and for two hours postprandially. Critically, we demonstrate that, unlike plasma acylated ghrelin, ghrelin autoantibody levels remained constant regardless of food intake status.

When considering the possible effects of ghrelin autoantibodies on the half-life of acylated ghrelin, the relative levels of both acylated ghrelin and ghrelin IgG should be considered. Acylated ghrelin levels decrease postprandially, while autoantibody levels remain constant. Therefore, the ratio of acylated ghrelin to ghrelin-reactive autoantibodies is reduced. In PWS, the constant elevation of ghrelin autoantibodies, which are believed to act as carrier proteins [5], could protect circulating ghrelin from degradation, potentiating its effect and contributing to hyperphagia, a key feature of this syndrome. Autoantibodies are thought to deliver acylated ghrelin to hypothalamic appetite-regulating centers, either directly or via the ghrelin receptor (GHSR) expressed by vagal afferents neurons. Pre-incubation of plasma with supraphysiological levels of UAG reduced autoantibody levels detected in plasma *ex vivo* in both the PWS and control groups, indicating that autoantibodies bind multiple ghrelin isoforms and, that in excess, UAG may compete with acylated ghrelin for binding. Kinetic studies have yet to be performed, however, it is possible that PWS patients exhibit distinct ghrelin-reactive antibody binding sites. Collectively, our data further implicate alterations of the ghrelin axis in PWS, and suggest that ghrelin autoantibodies can be targeted using non-orexigenic forms of ghrelin, thereby providing a novel therapeutic target for PWS and other ghrelin-associated disorders.

## Acknowledgements

The authors thank the study participants and their families and also acknowledge and thank Ms Sinead Archbold, Registered Nurse, for blood collection and clinical care of participants.

## Author contributions

Ms Crisp and Dr Nyunt are co-first authors, each with equal contributions to the manuscript. Drs Harris and Jeffery are co-senior authors. The project was conceived and designed by PLJ, ON, MH, IS, LKC and GC. Subject recruitment was performed by ON. GC and PLJ performed laboratory work. The manuscript was drafted by GC, PLJ, IS and LKC. All authors edited the final manuscript.

## Conflict of Interest Disclosures

None reported.

## Funding/Support

This research was supported by a grant to GC, LKC, IS, MH and PLJ from the Foundation for Prader-Willi Research (FPWR), and by an Australian Paediatric Endocrinology Group (APEG) grant to ON, PLJ and MH. IS is supported by a QUT Vice-Chancellor’s Senior Research Fellowship.

## Role of the Funder/Sponsor

The funding bodies had no role in the design and conduct of the study; collection, management, analysis, and interpretation of the data; preparation, review, or approval of the manuscript; and decision to submit the manuscript for publication.

## References

1. Cassidy SB, Schwartz S, Miller JL, Driscoll DJ. Prader-Willi syndrome. Genet Med. 2012;14(1): 10–26.

2. Haqq AM, et al. Serum ghrelin levels are inversely correlated with body mass index, age, and insulin concentrations in normal children and are markedly increased in Prader-Willi syndrome. J Clin Endocrinol Metab. 2003;88(1):174–8.

3. DelParigi A, et al. High circulating ghrelin: a potential cause for hyperphagia and obesity in Prader-Willi syndrome. J Clin Endocrinol Metab. 2002;87(12):5461–4.

4. Cummings DE, et al. Elevated plasma ghrelin levels in Prader Willi syndrome. Nat Med. 2002;8(7):643–4.

5. Takagi K, et al. Anti-ghrelin immunoglobulins modulate ghrelin stability and its orexigenic effect in obese mice and humans. Nat Commun. 2013;4:2685.

6. Fetissov SO. Neuropeptide autoantibodies assay. Methods Mol Biol. 2011;789:295–302.

